# Reconstructing Genotypes in Private Genomic Databases from Genetic Risk Scores

**DOI:** 10.1101/2020.01.15.907808

**Authors:** Brooks Paige, James Bell, Aurélien Bellet, Adrià Gascón, Daphne Ezer

**Affiliations:** The Alan Turing Institute, London, UK; Department of Computer Science, UCL, London, UK; Inria, France; University of Warwick, Coventry, UK; Department of Biology, University of York, York, UK

**Keywords:** Genomic privacy, Genetic risk scores, GWAS

## Abstract

Some organisations like 23andMe and the UK Biobank have large genomic databases that they re-use for multiple different genome-wide association studies (GWAS). Even research studies that compile smaller genomic databases often utilise these databases to investigate many related traits. It is common for the study to report a genetic risk score (GRS) model for each trait within the publication. Here we show that under some circumstances, these GRS models can be used to recover the genetic variants of individuals in these genomic databases—a reconstruction attack. In particular, if two GRS models are trained using a largely overlapping set of participants, then it is often possible to determine the genotype for each of the individuals who were used to train one GRS model, but not the other. We demonstrate this theoretically and experimentally by analysing the Cornell Dog Genome database. The accuracy of our reconstruction attack depends on how accurately we can estimate the rate of co-occurrence of pairs of SNPs within the private database, so if this aggregate information is ever released, it would drastically reduce the security of a private genomic database. Caution should be applied when using the same database for multiple analysis, especially when a small number of individuals are included or excluded from one part of the study.

## 1 Introduction

In a survey of genomic privacy experts, the long-term privacy of genomic information was deemed both the most important and the most challenging problem to overcome [11]. If an individual’s password or ID number gets leaked, then it is always possible to change it. However, it is impossible for a person to change their genetic code and they will pass part of it onto their children, so any information leaks can have long-term impacts on both the individual and their descendants. While much of the research focus on long-term privacy of genomic databases rests on the longevity of the encryption scheme [7], it is also important to remember that these genomic databases are not just sitting on a server somewhere, but are being continually utilised for making new scientific discoveries. Each time these databases are accessed and the scientific results are published, there is a risk that information will be leaked and that eventually this would enable an attacker to reconstruct private information held in the database.

Genomic researchers are already aware that some forms of aggregate data from their databases should not be released publicly, because there is a risk that an attacker may be able to determine whether a particular individual is a member of the database (a membership inference attack). For instance, such attacks have already been developed for summary statistics about the frequency of single nucleotide polymorphisms (SNPs) [2, 5, 15]. Membership inference attacks have also been developed for the case where a person is allowed to repeatedly query a database to learn if at least one individual contains a particular SNP [13, 14, 16]. These kinds of aggregate statistics about the frequency or presence/absence of a particular SNP might be useful to release to the broader research community, but it is not an essential output of the research process. However, the main research findings — i.e. the SNPs associated with the trait of interest and their strength of association — are essential to publish since the entire purpose of these genomic research projects is to uncover the relationship between genetic variants and phenotypic traits. Moreover, knowledge of these SNPs can lead to new diagnosis procedures or new potential drug targets, so their release is important for the public interest [18]. Yet, even this information can potentially leak private information about individuals in the database. For instance, [8] found that information about individuals in a genomic database is leaked when studies publish whether each SNP is correlated or anti-correlated to the trait of interest. It is important to quantify how much information is leaked by publishing these research findings, so that scientists can make informed decisions about when to publish their results and whether it is worth risking the privacy of the participants.

In this manuscript, we demonstrate that the kind of research output that is published from genome-wide association studies (GWAS) has the potential to leak enough information to recover the SNPs of individuals in the database (a reconstruction attack), under specific circumstances. In particular, we focus on the release of Genetic Risk Scores (GRS), a common research output for finding genetic associations with continuous traits [1, 3, 4, 10, 12, 20]. We also focus on cases where a database is repeatedly used to perform a GWAS analysis, but not all the individuals are part of all the analyses. This could be the case because some individuals drop out of the study or skip specific survey questions. Alternatively, some databases, such as 23andMe, may grow in size over time and allow several GWAS to be performed within a short period of time. Under these circumstances, we demonstrate that it is possible to completely reconstruct the SNPs of an individual using a custom Expectation-Maximisation (EM) algorithm. We also provide suggestions for avoiding this kind of attack.

To be clear, this manuscript focuses on the simpler case in which the exact same trait is investigated in multiple GWAS studies; however, we expect that some version of this attack may be developed in the near future for the case of multiple highly correlated traits.

### 1.1 Overview of scenarios that will be investigated

We demonstrate a series of reconstruction attacks that enable us to infer the genotypes of individuals in private genomic databases, based on publicly released GRS. These attacks will initially be deployed on a very favorable scenario, but the scope of the attack will be subsequently expanded, building up to the scenario shown in Figure 1. It is worth noting that the reconstruction attacks that we will describe do not depend on (i) how the SNPs were initially filtered or (ii) how strongly they associate with the trait of interest.

**Fig. 1.**
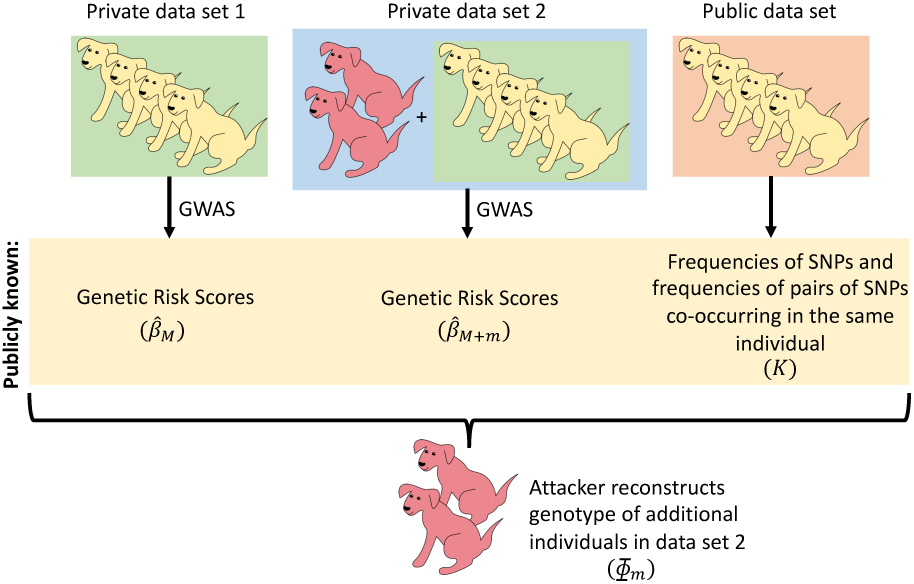
We investigate the case where two GWAS studies are performed on two data sets that mostly contain the same individuals. We reconstruct the genotype of those individuals added to the second study, using the GRS from each study and an estimate of SNP frequencies.

We will begin by investigating a simple scenario: two GWAS studies are performed to identify SNPs associated with the same trait, and the two studies use the same set of participants, except that the second study includes one extra individual. In addition, we will assume that we know the frequencies of each SNP and the frequencies that pairs of SNPs co-occur in the same individual. We assume that both studies publish the coefficients associated with the GRS models that they infer as part of the analysis.

Next, we will consider the case in which the second study includes more than one additional participant and demonstrate that in many circumstances this still allows us to easily reconstruct the individual genotypes of *all* the individuals that are found in the second study but not the first (see Section 3.2).

Afterwards, we will demonstrate that we do not need to know the precise frequencies of SNPs and frequencies of co-occurring SNPs, as long as we have a reasonable estimate of these values from public databases (see Section 3.3).

We also briefly discuss how loosening additional restrictions would impact our ability to predict individual genotypes. In particular, we analyse the case where the two sets of SNPs that are used by the two studies are not identical. These results imply that if two sets of GRS are released on two genetic data sets with largely overlapping populations, it may be possible to reconstruct the genotypes of those individuals who participated in one study but not the other (Figure 1).

## 2 Methods

Genetic risk score (GRS) models describe the relationship between a particular phenotype of interest and particular SNPs. These models are fit in a two-stage process: first, a reduced set of SNPs is selected from a potentially very large pool of candidates; then, this reduced set is used as the independent variables in a linear regression analysis. The set of SNPs is selected by first filtering for those that significantly correlate to the trait of interest, after controlling for other covariates. These SNPs are then further filtered to ensure that they are far apart from one another, in order to decrease the correlation between them.

In this setting, we suppose *M* individuals have taken part in a study, and *N* SNPs have passed the filtration steps to be used in a linear model. Let *y_M_* be the vector of *M* real-valued phenotypes, and *X_M_* be an *M* × *N* binary matrix, where *X_M_* [*i, j*] = 1 if individual *i* has SNP *j*. To include an intercept term in the linear model, we define a design matrix *Φ*_*M*_ to be the *M* × (*N* + 1) matrix

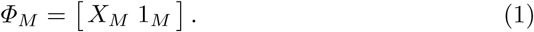

The GRS model parameter *β*_*M*_ is just the coefficient vector of the linear model

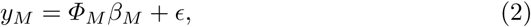

where *ϵ* is independent Gaussian noise. Given *Φ*_*M*_ and phenotypes *y_M_*, the maximum likelihood estimate of this parameter has a closed form

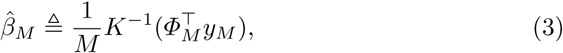

where we have defined the symmetric (*N* + 1) × (*N* + 1) matrix *K* as

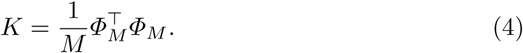

Now, suppose a second study is run, targeting the same phenotype, which adds a single extra individual with SNPs represented by the *N* length vector *x*_0_. This corresponds to adding the row 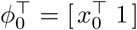 to the design matrix, and extending *y* with the additional phenotypic value *y*_0_ for the new individual. The updated estimator (i.e. the GRS values for the second study) is given by

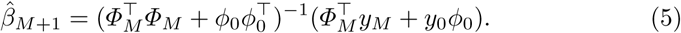

We assume that both GRS models 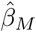 and 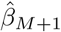 are released publicly. An attacker aims to use this knowledge to reconstruct *ϕ*_0_ (the genotype of the added individual). Through algebraic re-arrangement we find that:

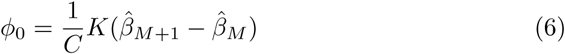

where *C* is a scalar, specifically 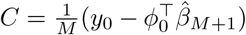. This means that *ϕ*_0_ is a scalar multiple of 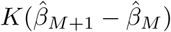.

Our approach thus centers on the use of the vector we define as *d*_1_,

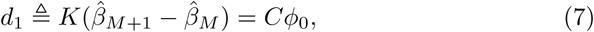

corresponding to a rescaled copy of the input SNP data in the design matrix *ϕ*_0_, which can be easily computed from the two parameter vectors if the matrix *K* is known. As we will see in Section 3.1, we can use *d*_1_ to *exactly* reconstruct the added individual with 100% accuracy.

We additionally consider the case where *m* additional individuals have been included in the second study, yielding a new GRS model 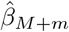 including these *M* + *m* participants. The extra rows of the design matrix now form a matrix *Φ*_*m*_ of size *m* × (*N* + 1), where each row is an individual that was added to the second study and each column is a SNP (and the last column contains only 1). The corresponding analog to Eq. (7) for multiple individuals, which we derive in Appendix B, is

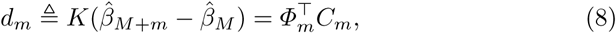

where *C*_*m*_ is a vector of length *m*. For sufficiently small *m* (relative to *N*), exact reconstruction of all *m* added individual genomes is also possible in this setting, following the algorithm we will introduce in Section 3.2.

The previous examples have focused on cases in which the participants in the first study are a subset of the individuals in the second study. In Appendix C we consider the case in which the first study has some participants that are not found in the second study and vice versa. We show that the same strategies for reconstructing the genome can be used as in the previous scenario that we discussed, in which multiple participants are added to the second study.

### 2.1 Estimation of *K*

As it turns out, the entries of matrix *K* correspond to simple population-level statistics of the SNPs, which could either be inadvertently released (under the assumption they would be safe to share), or could be estimated from another sample from the same population. In fact, the entries of *K* depend only on the SNP frequencies and SNP co-occurrence frequencies in the dataset:

− For *i* = 1, …, *N*: *K*_*ii*_ estimates the probability that SNP *i* has value 1 (i.e. the frequency of the SNP in the population).
− For *i* = 1, …, *N* − 1 and *j* > *i*: *K*_*ij*_ = *K*_*ji*_ estimates the probability that SNP *i* and SNP *j* are both 1 simultaneously (i.e. the frequency of SNP *i* and SNP *j* co-occurring in the same individual).
− For *i* = 1, …, *N* and *j* = *N* + 1: *K*_*ij*_ = *K*_*ji*_ also estimates probability that SNP *i* has value 1, i.e. *K*_*i,N*+1_ = *K*_*N*+1,*i*_ = *K*_*ii*_.
− Finally, *K*_*N*+1,*N*+1_ = 1.

Thus, knowledge of SNP frequencies and pairwise co-frequencies from the original study are all that is required in order to compute *K*. In the following Sections 3.1 and 3.2, we consider adding one and multiple individuals at once, respectively, in the setting where this matrix *K* can be estimated exactly.

However, while 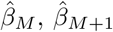 and *M* are likely to be published along with the study, an attacker would often need to estimate *K* from other publicly available data. Most studies will report some information about the study population (such as whether the study focused on individuals from a specific continent), which can help with estimating *K*. From this information, we can estimate the value of *K* in similar populations as those used in the study using publicly available data, e.g. from the HapMap project. Our additional experiments in Section 3.3 use a custom EM algorithm to find maximum likelihood estimates of *ϕ*_0_ when the matrix 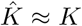 is estimated from independent public data. The derivation of this EM algorithm is given in Appendix D.3, and a formal analysis of the reconstruction error of *ϕ*_0_ given the error in 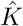 is found in Appendix D.1.

## 3 Results

The key observation from the previous section is that the vectors *d*_1_ and *d*_*m*_, derived from the change in parameter vectors 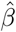 from a first study to a second study, *take only a finite number of values* thanks to the fact that the design matrices *Φ* contain only zeros and ones. In particular, when *m* new individuals are added to the second study, each entry of the vector *d*_*m*_ can only take at most 2^*m*^ values, and a zero value corresponds to the setting where all individuals have the most common variant for that SNP.

This section describes algorithmically how these vectors can be used to recover the genomes of the additional individuals, as well as empirical tests which use the Cornell Dog Genome dataset as a case study [6]. More details on the experimental setup can be found in Appendix A.

### 3.1 Complete reconstruction of one individual’s genotype when SNP frequency information is known

The first, most straightforward case is when only one participant is added between the first and second studies, i.e. where 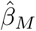 is the GRS for the first study (containing *M* participants), and 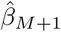 is the GRS for the second study as described in Eq. (3) and (5). Both of these are vectors of length *N* + 1, where the first *N* indices correspond to the relationship between each SNP and the trait and the last element is the intercept of the linear model. For now, we also assume we are in the setting where the matrix *K* is known, e.g. because the SNP frequency information has been publicly released.

Given *K*, 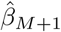 and 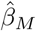, we can use *d*_1_ (a vector of length *N* + 1) to precisely determine the genotype of the individual who was added to the database. For each *i* = 1, …, *N*, the *i*^*th*^ entry of *d*_1_ is either equal to 0 if *ϕ*_0_ contains a 0 (i.e. the individual does not have the SNP at that index) or to *C* if *ϕ*_0_ contains a 1 (i.e. the individual has the SNP at that index). In other words, it is possible to *exactly* read off the SNPs of the added individual in this setting. Indeed, we tested this strategy on the Cornell Dog Database and found that we were able to reconstruct the genotype of the dog that was added to the second study with 100% accuracy, both on common and uncommon SNPs (see Figure 2(A)).

**Fig. 2.**
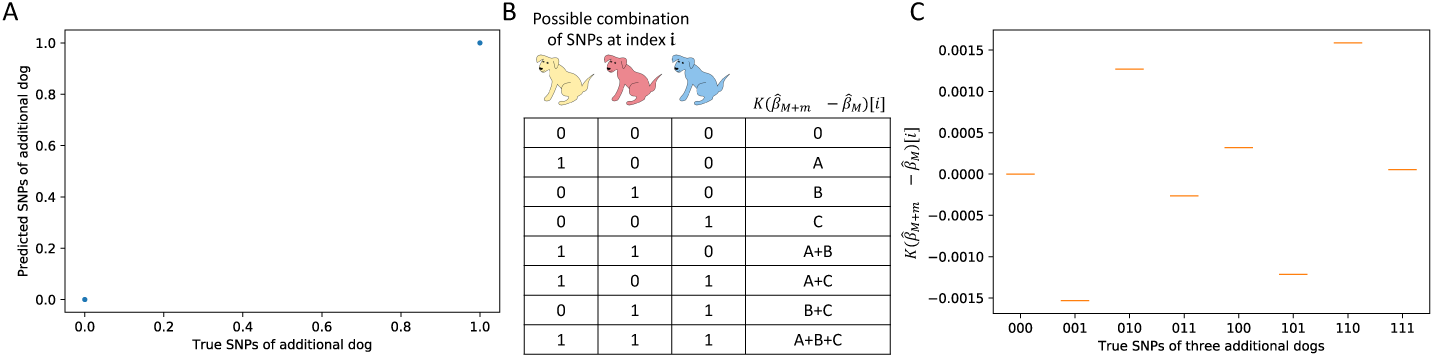
(A) We have perfect accuracy in reconstructing the genotype when *K* is known (using 200 random SNPs to estimate average breed weight in the Cornell Dog Database). (B) We can reconstruct all the genotypes of multiple dogs that are added to the second study and (C) this works in practice using the data from the Cornell Dog Database, as in (A).

### 3.2 Complete reconstruction of multiple individuals’ genotype when SNP frequency information is known

We now consider the case where *m* additional individuals have been included in the second study, yielding a new GRS model 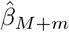 including these *M* + *m* participants.

Consider again Eq. (8) above. The *i*^*th*^ row of *Φ*_*m*_ is a binary vector that represents the combination of the *m* individuals who have SNP *i*. This means that, for a fixed value of *C*_*m*_, the value of the vector *d*_*m*_ at index *i* is uniquely determined by the combination of individuals who have SNP *i* (Figure 2(B)). In other words, there will be at most 2^*m*^ unique values taken by entries of *d*_*m*_, each corresponding to a combination of the values in vector *C*_*m*_ (see Figure 2(C)).

If we were to learn which values of *d*_*m*_ are also found in *C*_*m*_, then we could infer the complete genotypes of all the *m* individuals added to the second study. We would be able to reconstruct *m* complete genotype vectors, although it would be impossible to know which of the genotypes corresponded to which of the *m* individuals. In fact, in many cases it is extremely straightforward to determine which values in *d*_*m*_ correspond to values in *C*_*m*_. Here we describe a simple algorithm for finding *C*_*m*_ when there are exactly 2^*m*^ unique values in *d*_*m*_. If this is not the case, please see the more complete algorithm in Section G.

1. First, extract all unique, non-zero values from *d*_*m*_.
2. Find the sum of all pairs of values in this vector.
3. Find all values that are in (1), but not in (2). The values of *C*_*m*_ appear in this list. There is no way to know which value of *C*_*m*_ corresponds to which index, so for simplicity we can randomly assign them indices.
4. Each value in *C*_*m*_ corresponds to a specific individual who was added to the second study. Each value in *d*_*m*_ can be described as a sum of a unique combination of values in *C*_*m*_. For instance, if *d*_*m*_[*i*] = *C*_*m*_[*j*] + *C*_*m*_[*k*], this means that the SNP at position *i* is found in individual *j* and *k*, but no one else.

We tested this approach using the Cornell Dog Database, in a test scenario where the second study added three different dogs. We were able to uniquely identify the genotypes of all three dogs with 100% accuracy, both with common and uncommon SNPs (Figure 2(C)).

### 3.3 Accurate estimation of an individual’s genotype when SNP frequency information is estimated from a public database

Previously, we assumed that the attacker had access to the matrix *K*, which consists of population-level statistics on frequencies and co-occurrence frequencies of SNPs. While this could be released voluntarily by organisations which are not aware of the risk, we now consider the case where *K* is not directly available to the attacker but is instead estimated from a separate public database assumed to correspond to individuals from the same population.

We simulated this scenario using the Cornell Dog Database by taking one random set of dogs for building the GRS model, and a second non-overlapping set of dogs for estimating 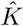. We compared the value of 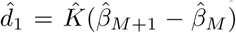 with the known value of *ϕ*_0_. We observe that 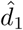 has significantly different values at indices where *ϕ*_0_[*i*] = 0 and *ϕ*_0_[*i*] = 1; examples for the cases where one and three dogs are added can be seen in Figure 3.

**Fig. 3.**
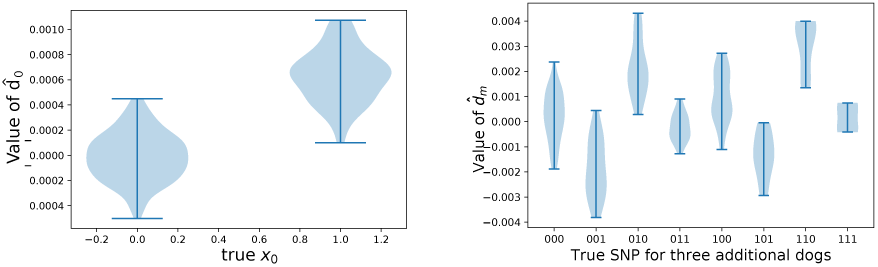
Example values taken by the noisy vector 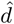, given the true value of the corresponding SNP in the genome. (Left) adding one new participant; (right) adding three new participants. These figures are analogous to those in Figure 2, albeit in the case where *K* is not known and instead estimated from an independent public database.

The main challenge is that the vector 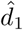 now includes additional noise, so we cannot simply use its entry at index *N* + 1 to estimate *C*, nor do the entries *i* with *ϕ*_0_[*i*] = 0 also correspond directly to 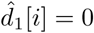. Instead, we develop a custom expectation-maximisation algorithm to find a maximum likelihood estimate of the constant *C* and recover *ϕ*_0_, i.e. to determine the probability that each *ϕ*_0_[*i*] = 0 or *ϕ*_0_[*i*] = 1, based on the value of 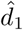 (see Section D.3 for details). We find that this method can successfully reconstruct the correct value of *ϕ*_0_[*i*] much better than a baseline which uses the public dataset to independently estimate the most common variant for each SNPs (see Figure 4). Crucially, we show that our approach is able to reconstruct, with relatively high accuracy, the genotypes of dogs even when they differ significantly from those in the public dataset (see Figure 5). This shows that our attack is able to extract information about the particular individuals that differ across the two studies, not merely about the general population as in the most-common-variant baseline. By definition, dogs that have genotypes that differ significantly from the general population have a higher proportion of uncommon SNPs, and the ability to recover these uncommon SNPs is particularly important from a privacy perspective. Indeed, uncommon SNPs can be used to identify a particular individual and are also more likely to be associated with disease phenotypes, which is sensitive information. In general, we find that the larger the public dataset available, and the more similar the dataset is to the unknown private dataset, the better we are able to reconstruct the genome of the added individual. Full details and description of the experimental setting are given in Section A. We also derive theoretical error bounds for our estimate of *ϕ*_0_ based on the error in 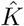 in Section D.1.

**Fig. 4.**
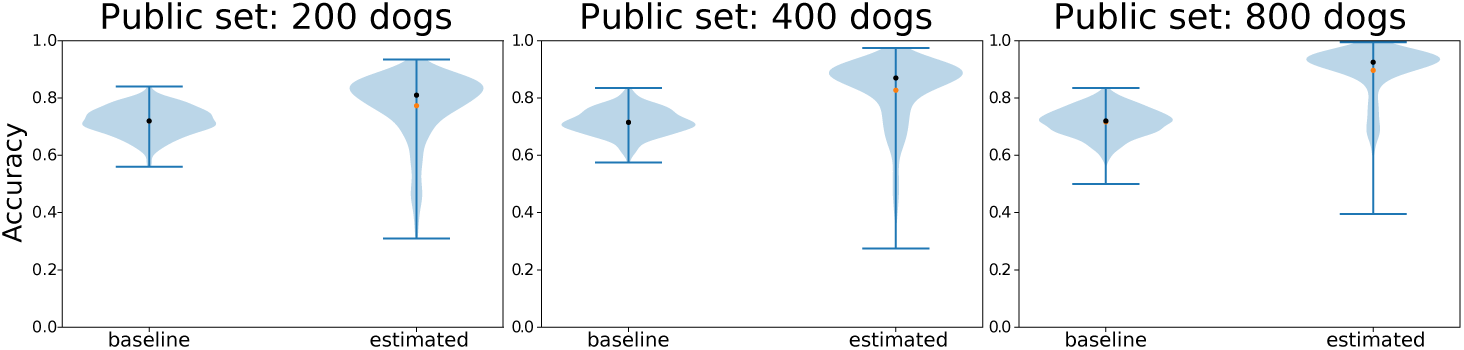
Accuracy at reconstruction of genomes *x*_0_ using EM estimation and a noisy estimate 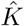, as compared to a natural baseline which always predicts the most common variant at each SNP locus. We use this as a baseline, because without any additional information about *β*_*M*_ and *β* _*M* +1_, the most accurate prediction of the dog’s genotype would be to predict the most common variant at each locus. Here we define accuracy as the proportion of SNPs that are correctly identified in the dog that was found in the second GWAS study, but not the first. Each distribution is constructed from 500 experimental test points, in which we (i) took 10 random splits of the full dog data set, assigning dogs to either the public and private data set (ii) for each split, we tested the reconstruction 50 times, each time adding a different randomly sampled dog to the second GWAS study. The private dataset always has 1000 individuals; the public test dataset is of increasing size, improving performance.

**Fig. 5.**
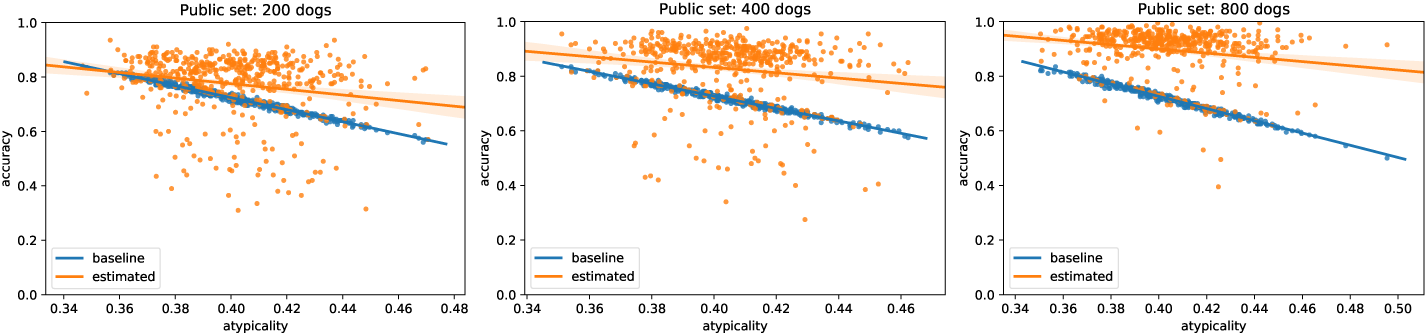
Results of Figure 4 broken down by individual dogs. Here each point represents a dog and we define *atypicality* as the proportion of *uncommon variants* that the dog has compared to the public database– for instance, if 51% or more of dogs in the public database have a *G* in a specific locus, but this dog has a *T*, then this would count towards the dog’s atypicality. In other words, dogs further to the right are less and less similar to average dog present in the public dataset (measured by percentage of different variants). In contrast to the most-common-variant baseline, our method generalizes well even to dogs which are highly dissimilar to those in the public dataset. Larger public databases (right) provide more accurate population estimates 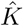, leading to more accurate reconstructions overall.

### 3.4 Accurate estimation of an individuals’ genotype when different SNPs are used in each study

When GRS models are constructed, the first step is to filter the set of SNPs down to a small set of SNPs that are (i) significantly correlated to the trait after covariates are considered and (ii) far apart from one another along the genome. If the two studies use two different sets of SNPs to construct the GRS model, it is still possible to recover whether or not each of the SNPs in the overlap is present in the new individual. This process is highly analogous to the previous cases and is detailed in Appendix F.

## 4 Discussion

In this manuscript, we demonstrate that private information is leaked when GRS models are published, specifically in the case where two sets of largely overlapping individuals are used for multiple studies. In particular, we show that we can recover SNPs from an individual in a private database—a reconstruction attack. Even though we would not have a *name* associated with this genotype, it may be possible to identify the individual once the genotypic data is available to the attacker. For instance, the attacker may have access to partial genotypic information of the individual and then be able to identify them. Alternatively, they could use the genotype information to predict ethnicity and other phenotypic traits that could then be used to uniquely identify the individual. We also note that even an incomplete reconstruction attack (in which only a proportion of the SNPs are correctly identified) is likely to be sufficient to perform a membership inference attack. Investigating the relationship between the reconstruction attack and the membership attack will be a subject of future research. Importantly, if the attackers were unable to link the genomic data with a particular individual, the reconstruction attack would still be a breach in privacy that could have serious consequences. For instance, the patient may have only consented to have their genomic data used in particular kinds of research studies, while the attacker may use the reconstructed genomic data for a different (potentially unethical) purpose.

### Suggestions for good practice

We provide a number of simple suggestions for good practice that would help limit this attack.

1. Aggregate statistics about the frequency of SNPs in the database or the frequency of co-occurrence of SNPs should never be released. We have shown that this information, combined with GRS, allows to precisely reconstruct individual genomes in various settings. It may be possible to release *noisy* versions of SNP frequency data, but this would be equivalent to releasing 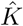 (our estimated *K* from the public database). With our EM algorithm, we have demonstrated that it is still possible to do some genotypic reconstruction with a noisy 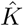, but this becomes harder as the noise in 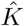 increases. However, providing a very noisy 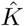 may be of limited utility to the scientific community.
2. If a genetic data set is intended to serve for multiple complementary analyses, it is important that all study participants are used in every analysis performed. If there is missing phenotypic data from a few individuals, they should not be included in *any* of the analyses performed, or their privacy may be compromised.
3. When multiple individuals are added in between two studies, then the ability to reconstruct the genomes depends on the number of SNPs being large relative to the number of individuals. In particular, if *m* new dogs are added, exact reconstruction is only possible using the approach in Section 3.2 if the number of SNPs *N* > 2^*m*^. Thus, we suggest to avoid releasing multiple studies which differ by fewer than log_2_ *N* individuals.

### Extensions and future work

While we have analyzed the case where the genome is represented by binary values of 0 or 1, often studies instead count the number of times each allele is present, which would lead to a design matrix *Φ* containing values 0, 1, or 2. In this scenario, *K* no longer contains the frequencies of SNPs and their co-occurrences, but something slightly more complicated that we describe in Appendix H. This does not dramatically change the approach in this paper, except in that the vector *d*_*m*_ can take 3^*m*^ possible values, rather than 2^*m*^. In practice, then, studies which use allele counts are somewhat more robust to attacks; the multiple dog reconstruction attack would likely be ambiguous if 3^*m*^ > *N*, rather than 2^*m*^ > *N*.

A possible countermeasure to our reconstruction attack could consist in randomly perturbing the GRS models before releasing them, as done in differentially private linear regression [19]. However, a naive application of this strategy could destroy the utility of the models. A formal and empirical analysis of the effectiveness of such protection against reconstruction attacks, as well as of the usefulness of the resulting GRS models to genomic researchers, is beyond the scope of this paper and left for future work.

Another countermeasure could consist in refraining from releasing precise information about the population structure of the study population to prevent the attacker from estimating *K* effectively. This would however limit the utility of the research study, because the researchers would not know to what populations the research applies to. An investigation of how differences in population structures impact *K*_*est*_ will be undertaken in public human genomic data sets in future work, but is also beyond the scope of this paper.

Our work has a number of limitations. For instance, we only test our EM algorithm on dog data. Dog populations may have different population structures than human populations due to selective breeding, so in the future we aim to test how properties of population structure will impact our ability to estimate *K* and the accuracy of our reconstruction attack. In addition, the expectation-maximisation algorithm described here is only explicitly described for the case in which one participant is added to the database, and the matrix *K* is estimated from a public database. We are currently extending the algorithm to the broader case in which multiple participants are added at once.

It may seem on the surface unlikely that two GWAS analyses will include nearly the same participants. One potentially common setting where this could arise is when a single study collects both genotype and phenotype data from a single set of participants, and releases multiple models to predict multiple traits. In this case, there may be a small number of individuals who are used in one analysis, but not the other; for instance, there may be a small subset of participants who skip a particular survey question that was used to collect phenotype information, and this is indeed evident in a recent study ([9]). In such settings, it could be very possible for multiple released GRS models to be computed on sets of individuals which differ by only a few participants. In future work, we aim to extend our analysis and attack to settings where multiple GRS models are released, each predicting different but highly correlated traits.

## A Experimental details

### Cornell Dog Database

To experimentally test the reconstruction attacks, we used data from the Cornell Dog Genome Database, which contains data about SNPs from a wide range of dog breeds and a number of associated phenotypic traits. The two traits we focused on were *average breed weight* and *average breed height*, because these two phenotypes had the fewest number of missing values. For the initial investigation, we binarised the genotype matrix—considering all heterogenous alleles to have a value of 1. (We also repeated the analysis with the original genotype matrix.) Only common SNPs (i.e. SNPs that were found in 25% to 75% of the dogs) were used, leaving 23,497 SNPs. For each linear model built, *M* = 1000 dogs were randomly sampled as the “private” dataset and *N* = 200 SNPs were randomly selected. To ensure that the SNPs that were sampled were spatially distributed, the SNPs were randomly sampled in a stratified way, so one SNP was selected in every 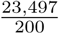-sized bin.

### Experiment with imprecise *K*

First, two linear models were constructed to predict average breed weights: one with the *M* = 1000 randomly sampled dogs and another that contained 1 additional randomly sampled dog. This gives 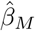 and 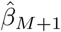. To mimic the process of estimating *K* from a public database, we randomly sampled an additional 200, 400, or 800 dogs that were not included as part of the original set and used this to estimate *K*, which we refer to as 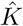. Now we could calculate 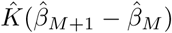 and compare this to the known *ϕ*_0_ for the additional dog from the second study. These additional dogs are taken from a third “test” dataset, disjoint from both the public and private data. The plots in Figures 4 and 5 are produced by re-running the algorithms across 10 random public / private / test splits, where the “test” dataset has 50 dogs which are each individually considered as candidates for the (*M* + 1)^th^ dog added to the private dataset.

## B Adding multiple dogs

Here we explain Equations (7) and (8). Note that the former is a special case of the latter so we will only explain the latter in detail. First note that by definition

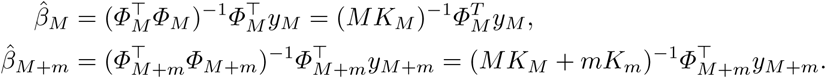

Substituting these into the left hand side of the following equation gives the right hand side:

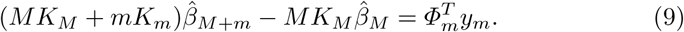

This equation can be rearranged to give

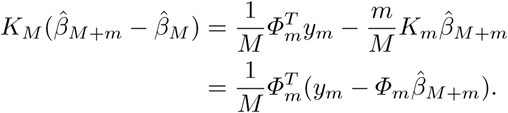

Defining the length *m* vector 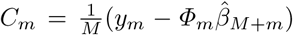 yields the form used in Equation (8). For the special case of *m* = 1, *C*_*m*_ is a scalar and we recover Eq. (7).

## C Case in which each GWAS study adds two new sets of participants

This manuscript mostly explores the case in which one study’s participants are a subset of the other study’s participants. Here we demonstrate that this is equivalent to the case where each of the two studies contain a small number of participants that are not found in the other study.

In particular, let us say that the first study has *M* + *a* participants and the second study has *M* + *b* participants, where the first *M* participants are shared between the studies, but there are *a* participants that are found in the first study but not the second, and *b* participants that are found in the second study but not the first. Following on from Equation 9, we see that:

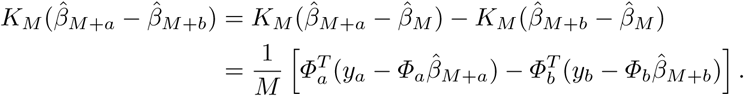

Let us define the following (*N* + 1) × (*a*+ *b*) matrix obtained by concatenating the two genotype matrices:

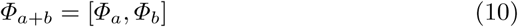

and the following *a* + *b* length vector:

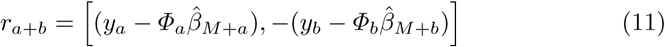

Then this gives us:

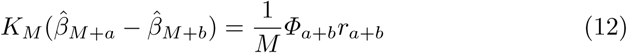

This means that having two non-overlapping participant sets is equivalent to the setting in which the first study is a subset of the second (only *m* is now *a* + *b*).

## D Estimating *K*

If the true matrix *K* is unknown, it can be estimated with public data. We denote this estimator by 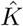. In order for 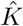 to be an accurate estimate the data that it is generated from must be drawn from the same (or a sufficiently similar) population as that used in the private study. We will model this assuming no discrepancy between population distributions, however when we discuss how to evaluate whether the estimate is good that assessment should account for this systematic error as well.

In the following we are primarily concerned with the error due to the subsampling in both the private and public data sets. For convenience we only consider the case of adding a single individual, though the generalization is quite straight-forward.

### D.1 Analytic bound on 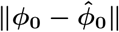

If 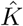 is substituted for *K* in our reconstruction equation (7) we get an approximation of *ϕ*_0_ which we denote 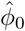. We would like to bound the (relative) error between *ϕ*_0_ and 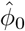. In the following, we ignore the constant factors *C* and *Ĉ* for simplicity, noting that these scaling factors are estimated from the resulting *ϕ*_0_ or 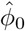 anyway. We thus consider 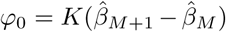 and 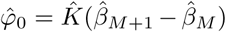. Using ‖· ‖ on vectors, and also on matrices to denote the corresponding operator norm. The relative error between *ϕ*_0_ and 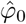 is given by:

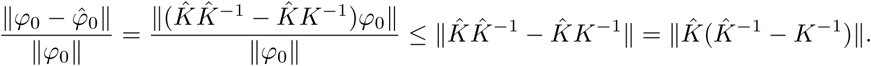

Note that 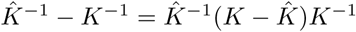 and hence

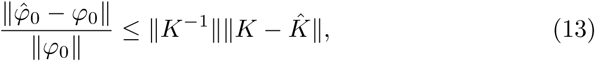

This means we can bound the error by two quantities. The term ‖ *K*^−1^‖ is bounded above by 1*/* min(eig(*K*)), which is finite as soon as *K* is non-singular. This is not a strong requirement as in the case of linear regression it is required for 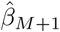 and 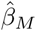 to exist. Note that in the case of L2-regularized linear regression (i.e., ridge regression), *K* is replaced by *K* + *λI* where *λ* is the regularization parameter, and we can directly bound this term by *λ*.

The key term in (13) is 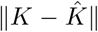, the error in estimating *K* by 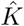. Let us assume that the public database used to obtain 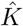 follows the same distribution as the private database used to fit the GRS models. Denote by 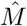 the number of individuals used to estimate 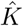. Then, under classic boundedness assumptions and leveraging matrix concentration inequalities such as matrix Bernstein [17] we can show that 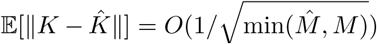. This shows that the error in estimating *K* is small as long as the private and public databases are large enough.

### D.2 Modelling the error in 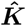

In this section we define a model to capture the error in 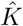, which leads to the expectation maximization algorithm for estimating *ϕ*_0_ which is used in the experiments.

As our estimated 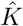 drifts from the true *K*, this expression 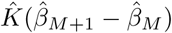 would produce a wider range of values than just 0 and *C*. Let *ϵ*_*ij*_ ∼ 𝒩 (0, *σ*^2^) be independent noise, which we assume corrupts each element of *K*_*ij*_; i.e. given the estimated matrix 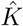, suppose

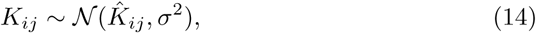

for some small *σ*^2^. This is clearly an oversimplification (as we know *K* is e.g. bounded and symmetric), but is a useful starting point that allows derivation of a simple estimation algorithm. For notational brevity, in this and the following section we define the vector

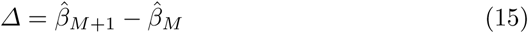

which corresponds to the difference between the two GRS model parameter vectors. Given the true value of *K*, the system of equations

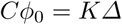

relates the known quantity Δ and the Gaussian-distributed *K* with the unknown value of *C* and of the vector *ϕ*_0_.

If *K* is Gaussian (following Eq. (14)), then the linear transformation *K*Δ is Gaussian as well. We denote each of the rows of *K* as a vector *k*_*i*_, *i* = 1, …, *N*; then for each row, the scalar value

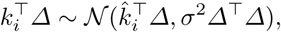

meaning overall the vector *K*Δ is distributed 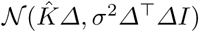. With some algebraic re-arrangement, and since for the true underlying value of *K* we have *K*Δ = *Cϕ*_0_, we can write this as

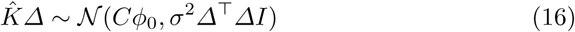

where *C* and *σ* are parameters we need to estimate. The vector 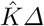 is observed “data”, computed from the public SNP database and the two released parameter vectors. We can model each of the entries of *ϕ*_0_, which are zeros and ones, as Bernoulli distributions, whose prior probabilities correspond to the public dataset estimated frequencies. This suggests a model for 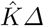 which is akin to a constrained mixture of Gaussians.

### D.3 Derivation of EM algorithm for reconstructing an individual’s genotype

Given the model in Eq. (16), we can define an EM algorithm for maximum likelihood estimation of *C* and *σ*^2^, which then permits easy inference for each entry of *ϕ*_0_.

Denote the entries of *ϕ*_0_ as *z*_1_, …, *z*_*N*+1_; denote their prior probabilities as *α*_1_, …, *α*_*N*+1_, where *α*_1_, …, *α*_*N*_ are the (public) population frequencies for each SNP, and *α*_*N*+1_ = 1.0. Let *x*_1_, …, *x_M_* denote the entries of the fixed (observed) vector 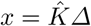, which is distributed

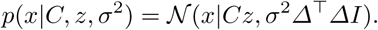

Supposing we know values of *C, σ*^2^, to estimate the entries of *ϕ*_0_ we want to find *p*(*z*|*x, C, σ*^2^),

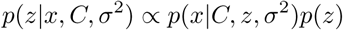

For each *z*_*i*_, *i* = 1, …, *N*, we define the quantity

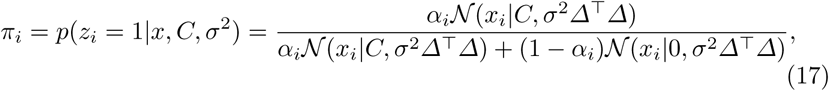

the posterior probability of each particular entry taking a value of 1, rather than 0. A maximum likelihood algorithm to estimate *C, σ*^2^ proceeds by alternately:

1. Given *C, σ*^2^, estimate the posterior distribution *π* = *p*(*z*|*x, C, σ*^2^);
2. Given the posterior *π*, maximize ℒ = *E*_*π*_[log *p*(*x*|*C, z, σ*^2^)] with respect to *C* and *σ*^2^.

To maximize *C* and *σ*^2^, we first compute the derivatives of

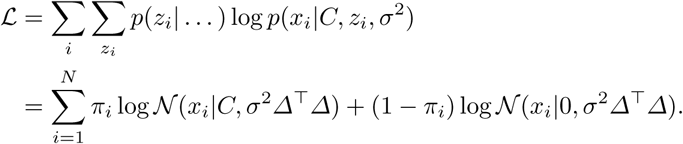

We take derivative w.r.t. *C*,

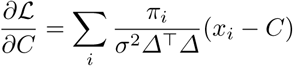

and set equal to zero to find

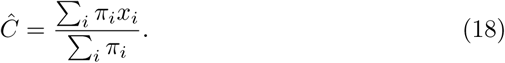

For the noise term *σ*^2^, we have

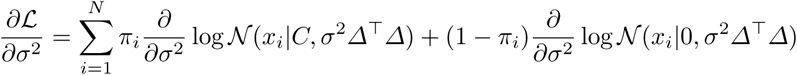

which with some algebraic rearrangement becomes

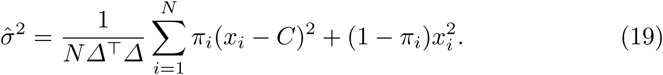

These updates taken together can be used to define an EM algorithm which optimizes the values of *C* and *σ*^2^, despite the fact that the entries of *ϕ*_0_ are unknown; once *C* and *σ*^2^ are then known, the vector *π* will give probability estimates for each entry of *ϕ*_0_.

The overall EM algorithm can be summarized by the following iterative updates:

1. 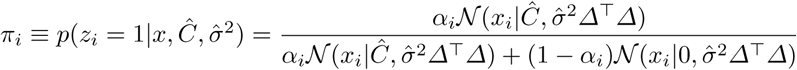,
2. 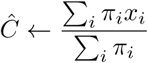,
3. 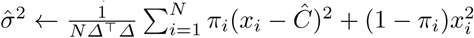.

To initialize the algorithm, we can set *π*_*i*_ to some initial probabilities, and find initial values for 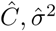; we experimented with both setting to the prior probabilities per-SNP estimated from the public data, as well as to the vector of all zeros (corresponding to a “hard” initialization at the value of the baseline estimate), and found no qualitative difference in performance.

## E Scaling of EM algorithm with size of private dataset

Figure 6 demonstrates the change in accuracy of the EM algorithm over a range of different private database sizes. For this test, a synthetic dataset with 100 SNPs and 1,000,000 individuals is generated; 10,000 are held out as a public database, and 30 individuals are taken as a fixed test dataset of new dogs to add and are used to estimate EM algorithm accuracy, across increasingly large private database sizes. The algorithm has stable performance for increasingly large private databases.

**Fig. 6.**
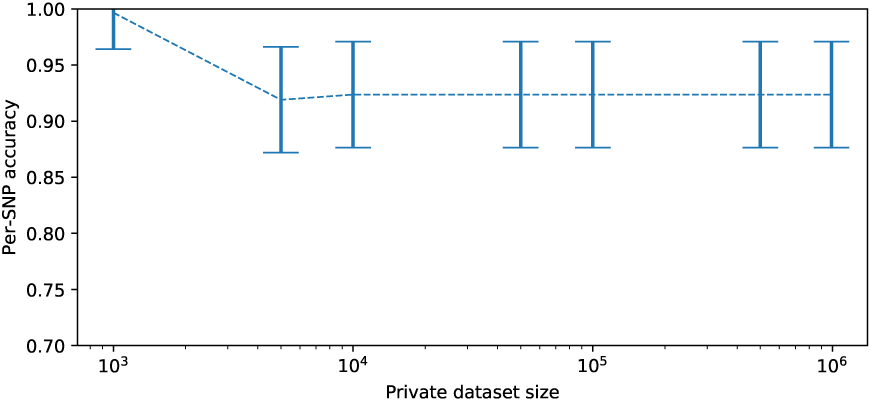
Accuracy at reconstruction of the genome of one additional individual, using EM estimation and a noisy estimate 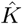, measured as the size of the initial private database increases. For very small private databases, accuracy is very high, as changes in entries of *β* are clearly attributable to the new individual. Beyond a certain threshold, overall accuracy is quite stable. Error bars show mean and two standard deviations.

## F Estimating *ϕ*_0_ with different SNP sets

Here we analyse what can still be said in the event that the two studies do not use exactly the same set of SNPs. We will still assume the sets of SNPs considered to have a significant overlap.

For this purpose we will need a greater variety of notation. A primed variable denotes that it corresponds to the second set of SNPs, e.g. *K*′ is the co-occurrence matrix from the original *M* users for the second experiment. If a vector or matrix is surrounded by square brackets this denotes that object but with any rows or columns corresponding to SNPs not in the overlap removed, e.g. [*K*] denotes the co-occurrence matrix from the first experiment restricted to the overlapping SNPs.

As before, from the first experiment, we have

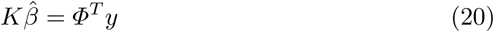

and now, from the second experiment, we have

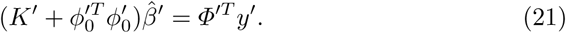

Taking the difference between these expressions, as before, gives

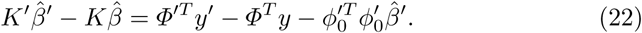

Restricting to the overlapping set gives that

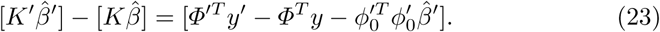

Noting that [*K*] = [*K*′] and that 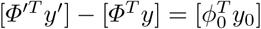 we get that

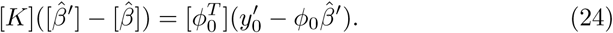

Analogously to the previous cases 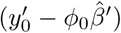 is a scalar which we can label *C* and we get

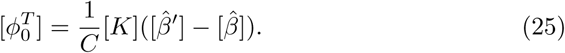

Thus if *K* is known it can be used to deduce whether the additional individual has each of the SNPs in the overlapping set. If *K* is not known exactly it can be estimated from public data just as in the same SNP case.

## G Algorithm for identifying unique genotypes of multiple dogs, when a precise *K* is known

While the simple approach described in the main manuscript will work in many cases, there are a few special circumstances where a more complex algorithm may be required. In particular, it would not work if there are combinations of SNPs that are not observed among the individuals added to the database. For instance, if there is not a single SNP location where the first individual has a SNP variant and the others do not, then we would miss the corresponding value in *C*_*m*_. However, it is still possible to identify all the values in *C*_*m*_ through a more complex algorithm:

1. First, extract all unique, non-zero values from *d*_*m*_.
2. Find the sum of all pairs of values in (1).
3. Find all values that are in (1), but not in (2).
4. If there are exactly *m* values in (3) and the sum of these values equal the last value of *d*_*m*_ (corresponding to the intercept term), then you have found the correct values of *C*_*m*_.
5. Otherwise, this suggests that there are one or more elements of *C*_*m*_ that are missing from (3) and possibly a few values in (3) that are not in *C*_*m*_.
6. Begin by subtracting every pair of values in (3). These are now also potential values of *C*_*m*_
7. Search for a set of *m* values from (3) and (6) that sum to the last element of *d*_*m*_. There may be more than one set of values for which this is true.
8. If this search is unsuccessful, repeat steps 6-7. Eventually, a set of *m* values summing to *d*_*m*_ should be found.
9. If more than one possible set of values is found for *C*_*m*_ in (7), it is still possible to compare these sets and identify which is the most likely to contain the true values of *C*_*m*_. For each possible *C*_*m*_ vector, a set of genotypes can be constructed for the *m* additional individuals. Using the frequencies of each SNP, it is possible to calculate the probability of observing each genotype. The set of values that produces the most likely genotypes for the *m* individuals is most likely to be the correct one.

Additionally, this algorithm depends on the fact that it is extremely unlikely that if someone were to sample three random continuous numbers *i, j* and *k*, it would just so happen that *i* + *j* = *k*. There is an extremely small chance that a value of *C*_*m*_ would be un-discoverable because of a coincidence of this nature.

## H Description of *K* when the genotypes are non-binary

In many cases, GRS are calculated on genotype matrices that are non-binary. In particular, they may take on three discrete values 0, 1 and 2, where 0 indicates that the most common variant is homozygous, 1 indicates that the individual is heterozygous for the uncommon variant, and 2 indicates that the individual is homozygous for the uncommon variant.

If this is the case, the description of *K* will change. However, it is still the case that the entries of *K* depend only on the SNP frequencies and SNP co-occurrence frequencies in the dataset, and that knowledge of SNP frequencies and pairwise co-frequencies from the original study, are all that is required in order to compute *K*.

– For *i* = 1, …, *N*: *K*_*ii*_ = *p*_*Aa*_ + 4*p_AA_* where *p*_*aa*_ is the frequency of individuals being heterozygous for the uncommon variant and *p_AA_* is the frequency of individuals being homozygous for the uncommon variant.
– For *i* = 1, …, *N* − 1 and *j* > *i*: *K*_*ij*_ = *K*_*ji*_ = *p*_*Aa/Bb*_ + 2*p*_*AA/Bb*_ + 4*p_AA/BB_* where *p*_*Aa/Bb*_ is the frequency that both SNPs are simultaneously heterogygous, *p*_*AA/Bb*_ is the frequency that one SNP is homozygous for the rare variant and the other is heterogygous simultaneously, and *p_AA/BB_* is the frequency that that uncommon variants are found to be homozygous simultaneously.
– For *i* = 1, …, *N* and *j* = *N* + 1: *K*_*ij*_ = *K*_*ji*_ = *p*_*Aa*_ + 2*p_AA_*.
– Finally, *K*_*N*+1,*N*+1_ = 1.

